# Myosin IIA motor regulates attaching-effacing bacteria interactions with intestinal epithelium

**DOI:** 10.1101/2025.10.11.681730

**Authors:** Nayden G. Naydenov, Atif Zafar, Susana Lechuga, Armando Marino-Melendez, John A. Hammer, Velia M. Fowler, Christine McDonald, Kenneth G. Campellone, Andrei I. Ivanov

**Author notes:** Correspondence: Andrei I. Ivanov, PhD. Department of Inflammation and Immunity Cleveland Clinic Research Cleveland, OH 44195, USA.

## Abstract

Attaching effacing (A/E) bacteria, such as Enteropathogenic *E. coli* (EPEC) and *Citrobacter rodentium*, colonize intestinal epithelial cells (IECs) by inducing remodeling of the epithelial cytoskeleton and formation of prominent actin pedestals at bacterial attachment sites. While non-muscle myosin II (NM II) is a key regulator of the actin cytoskeleton, whether it regulates IEC colonization by A/E pathogens is not known. To address this question, we targeted NM IIA and NM IIC, the NM II paralogs expressed in IECs. Our *in vivo* studies utilized mouse models with either intestinal epithelial-specific deletion of NM IIA (NM IIA cKO mice), expression of a NM IIA motor domain mutant, or total deletion of NM IIC (NM IIC tKO mice). *In vitro* experiments utilized IECs (HT-29cF8 and Caco-2BBE) with CRISPR-Cas9-mediated deletion of NM IIA or NM IIC. In addition, NM II activity *in vitro* was modulated pharmacologically, using either the pan-myosin inhibitor, blebbistatin, or a specific NM IIC activator, 4-hydroxyacetophenone (4-HAP). NM IIA cKO and NM IIA mutant mice demonstrated higher *C. rodentium* colonization along with more severe mucosal inflammation and colonic crypt hyperplasia as compared to their controls. By contrast, NM IIC tKO mice was indistinguishable from their control with regard to *C. rodentium* colonization. Blebbistatin treatment increased EPEC attachment to IECs monolayers, whereas 4-HAP did not affect bacterial attachment. Genetic knockout of NM IIA, but not NM IIC, increased EPEC adhesion to IEC monolayers. Importantly, the increase in EPEC attachment exhibited by NM IIA-deficient IECs required intact bacterial Type 3 secretion system and functional Tir effector, indicating that NM IIA functions in actin pedestal assembly. In summary, we describe a novel role for NM IIA in limiting intestinal epithelial colonization by A/E pathogens via inhibition of pathogen-induced remodeling of the actin cytoskeleton.

## Introduction

Remodeling of the actin cytoskeleton is a key mechanism utilized by different pathogenic microorganisms to colonize mammalian cells ^1, 2^. Pathogens perturb actin filament architecture and dynamics and trigger assembly of new cytoskeletal structures to support their adhesion, invasion, and dissemination within the host cells and tissues ^1, 2^. A group of enteric pathogenic bacteria is known to induce dramatic remodeling of the mammalian actin cytoskeleton by attaching to the host cell surface and injecting different effector proteins that elicit a variety of intracellular signaling events. Such pathogens include Enteropathogenic *Escherichia coli* (EPEC) and Enterohemorrhagic *Escherichia coli* (EHEC), known to cause serious diarrheal diseases in children and immunocompromised adults ^3, 4^. Similar features are also attributed to a mouse pathogen, *Citrobacter rodentium*, which is commonly used as an experimental model for EPEC and EHEC infection ^5, 6^. Collectively, these bacteria are referred to as ‘attaching/effacing’ (A/E) bacteria due to their ability to destroy microvilli and form attaching/effacing lesions at the apical surface of infected intestinal epithelial cells (IECs) ^7–9^. The hallmark of the A/E pathogen-induced remodeling of the host cytoskeleton is the assembly of actin-rich pedestals underneath the plasma membrane where attached bacteria reside ^1, 2, 10^. These pedestals represent the most vivid morphological alteration of host cells triggered by the attached pathogen, and they appear to be essential accelerators of bacterial infection ^11^. The ability of A/E pathogens to remodel the host cytoskeleton and build actin pedestals depends on crucial bacterial virulence factors, such as a type III secretion system (T3SS), the adhesin intimin, and its receptor Tir. The T3SS system represents a syringe-shaped multiprotein complex penetrating the host membrane that is used to inject many bacterial effector proteins into the host cells ^12–14^. Bacterial Tir and other secreted effectors stimulate actin filament polymerization within the vicinity of the attached bacteria, leading to actin pedestal assembly ^12–14^. The mechanisms by which the A/E pathogens subvert the host actin polymerization machinery have been extensively investigated over several decades ^10, 15–18^. However, the pedestals contain many other cytoskeletal modulators, including actin filament depolymerizing, filament cross-linking, membrane-tethering proteins as well as cytoskeletal motor proteins, myosins ^16, 17^. For most of these actin-binding proteins, their roles in pedestal assembly and host cell colonization by A/E pathogens remain poorly understood.

One of the most obvious knowledge gaps is lack of understanding of how non-muscle myosin II (NM II) motors control A/E pathogen interactions with mammalian cells, particularly in the intestinal epithelium. NM II is a critical regulator of virtually all actin-based cellular processes and possesses a dual ability to translocate and cross-link actin filaments ^19, 20^. IECs express two different NM II motors, NM IIA and NM IIC ^21, 22^. These motors are enriched at the apical pole and intercellular junctions of enterocytes and regulate epithelial adhesion and microvilli dynamics ^21–23^. Several lines of indirect evidence suggest that NM II may play role in the A/E pathogen-induced remodeling of the actin cytoskeleton. First, immunofluorescence labeling and mass spectroscopy analysis detected NM II in EPEC pedestals ^24, 25^. Second, EPEC effector proteins, EspB and Map, are known to interact with NM II ^26, 27^. Lastly, EPEC and *C. rodentium* have been shown to stimulate NM II activity by increasing phosphorylation of myosin light chains (MLC) ^28–31^. While such NM II activation is thought to be essential for disrupting apical junctions in EPEC and EHEC-infected intestinal epithelium ^30–32^, it remains unknown whether NM II controls pedestal assembly and bacterial attachment to host cells. This study was designed to fill this knowledge gap and examine the roles of NM II motors in regulating EPEC and *C. rodentium* interactions with the intestinal epithelium *in vitro* and *in vivo*. Our results highlight NM IIA as a unique negative regulator of A/E bacterial attachment to IECs, controlling bacteria-induced remodeling of the actin cytoskeleton and pedestal formation.

## Methods

### Antibodies and other reagents

The following primary polyclonal (pAb) and monoclonal (mAb) antibodies were used to detect cytoskeletal proteins and EPEC: NM IIA pAb, (BioLegend, San Diego, CA cat.# 909801, Anti LPS pAb (Abcam, Waltham, MA cat.# 35654), NM IIC (D4A7) cat.#8189S, and anti-GAPDH (14C10) mAbs (Cell Signaling, Beverly, MA, cat.# 2118); Alexa Fluor-488 or Alexa Fluor-568-conjugated donkey anti-rabbit or donkey anti-mouse secondary antibodies, Alexa Fluor-488, Alexa Fluor-555 and Alexa Fluor-647-labeled phalloidin were obtained from (ThermoFisher Scientific, Waltham, MA). Horseradish peroxidase-conjugated goat anti-rabbit and anti-mouse secondary antibodies were acquired from Bio-Rad Laboratories (Hercules, California). All other chemicals were obtained from Millipore-Sigma (St. Louis, MO).

### Animals

The mouse line with intestinal epithelial-specific knockout of NM IIA was established by crossing NM IIA flox animals with constitutive villin-Cre mice, as previously described^22^. Transgenic NM IIA mouse lines with replacement of mouse NM IIA gene with GFP-fused wild type human NM IIA or the R702C NM IIA mutant (referred to as NM IIA WT-GFP and NM IIA R702C-GFP, respectively) were described previously ^33, 34^ and were provided by Dr. Velia M. Fowler (University of Delaware). A mouse strain with total knockout of NM IIC, (C57BL/6J-Myh14, NCBI ID 71960) was generated by (Cyagen, Santa Clara, CA), and provided by Dr. John Hammer (National Institutes of Health). The animal colonies were maintained under pathogen-free conditions with a 12 h light/dark cycle in the Cleveland Clinic vivarium. Mice were fed a standard chow diet of Envigo Diet 2918 (irradiated) and provided with filtered and chlorinated municipal water *ad libitum*. All procedures were performed as Approved by Cleveland Clinic Institutional Animal Care and Use Committee (IACUC protocol number 00001872) per the National Institutes of Health Animal Care and Use Guidelines.

### Bacterial strains and in vitro infection experiments

*Citrobacter rodentium* strain DBS100 was obtained from the American Tissue and Cell Collection (ATCC, Manassas, VA. #51459). EPEC strain E2348/69 (Serotype O127:H6) was provided by Dr. Kaper for distribution by BEI Resources NIAID, NIH (Cat # NR-50518). EPEC mutant strains: EPEC-Δ*tir*, EPEC-Δ*tir*+pHA-Tir(WT), EPEC-Δ*tir*+pHA-Tir(Y454F/Y474F), and EPEC ΔT3SS were described previously ^35–38^.

Single colonies of bacteria were propagated overnight by culturing in the Luria-Bertani (LB) medium with agitation at 37°C. Subsequently, EPEC strains were subcultured for 3 h in the Dulbecco Minimum Essential Medium (DMEM) without antibiotics. Bacterial concentration was estimated by measuring optical density at 600 nm, and the suspensions were diluted in their growth media to the desired MOI. For the majority of in vitro experiments, epithelial cells were infected with EPEC at MOI 5:1 for 3 h. For immunolabeling experiments, EPEC infection was performed in confluent IEC monolayers growing on collagen-coated coverslips. For the CFU measurements, the infection was performed in confluent IEC monolayers growing on collagen-coated 24 well plates. *C. rodentium* infection in mice is described below.

### Cell lines and generation of CRISPR/Cas9-mediated NM II knockout IEC

Caco-2BBE (# CRL-2102) and Cos-7 (# CRL-1651) were purchased from ATCC. HT-29 cf8, a well-differentiated clone of HT-29 cells ^39, 40^ was provided by Dr. Judith M. Ball, (College of Veterinary and Biomedical Sciences, Texas A&M University, College Station, TX). HT-29, Caco-2, and Cos-7 cells were cultured in DMEM medium supplemented with 10% fetal bovine serum (FBS), HEPES, non-essential amino acids, and penicillin-streptomycin antibiotics. Knockout of NM IIA and NM IIC in IEC lines was achieved using the CRISPR/Cas9 V2 system. Single guide RNAs (sgRNAs) targeting *NM IIA* or *NM IIC* were designed via the CRISPR design tool developed by the Feng Zhang Lab (http://crispr.mit.edu, McGovern Institute, MIT, Boston, MA, USA). The sgRNA sequences used were: NMIIA sg1–ACGCCACGTACGCCAGATAC and NMIIA sg8– CTGAGTAGTAACGCTCCTTG and NM IIC sg1-CCTCGGTCACTGCGTACACG, and NM IIC sg3-GATTACTCACGTGCAGAGAA. Each sgRNA was cloned into the BbsI site of the lenti-CRISPR v2 vector (Addgene, Watertown, MA, #52961), with successful insertion confirmed by sequencing. Lentiviral particles were produced in HEK293T cells co-transfected with the lenti-CRISPR-sgRNA plasmids and packaging plasmids pCD/NL-BH*DDD (Addgene #17531) and pLTR-G (Addgene #17532), using the TransIT-293 transfection reagent (Mirus Bio, Madison, WI, USA). To establish stable knockouts, HT-29 and Caco-2 BBE cells were transduced with the resulting lentiviruses and subjected to puromycin selection (5 μg/mL) for 7 days. Control cells were generated by transduction with lentiviruses carrying a non-targeting sgRNA, followed by the same selection protocol. For bacterial infections, HT-29 and Caco-2 BBE cells were cultured for 7 and 14 days respectively to allow epithelial cell differentiation.

### Generation and expression of human NM IIA mutants

To generate NM II mutants, the backbone plasmid CMV-GFP-NMHC IIA, in which the human *NM IIA* gene is fused to GFP (Addgene, #11347), was used. Individual point mutations N93K and R702C were introduced using the QuikChange Lightning Site-Directed Mutagenesis Kit (Agilent Technologies, Santa Clara, CA, # 200515). Phosphorylated primers GCACCGAGGCTTCCTTGAGGCACGTGA (N93K) and CTGGCGGCAGATACAGATGCCCTCGAGAA (R702C) were used in the PCR-based mutagenesis reaction, performed according to the manufacturer’s instructions using a C1000 Touch Thermal Cycler (Bio-Rad laboratories). The PCR products were digested with the restriction enzyme DpnI to remove methylated and hemi-methylated parental DNA templates. The resulting reaction mixtures were transformed into XL10-Gold® ultracompetent cells. Plasmid DNA was isolated from individual colonies using the PureYield Miniprep System (Promega, Madison, WI, #A1222). The presence of the desired mutations was verified by DNA sequencing. For transfection Cos-7 cells were seeded onto collagen-coated coverslips at 50% confluence and transfected the following day with the indicated plasmids at a final concentration of 0.5 µg/mL using the TransIT-2020 transfection reagent (Mirus Bio, Madison, WI, #MIR 5404) according to the manufacturer’s instructions. Forty-eight hours post-transfection, cells were infected with EPEC at MOI 5:1 for 3h. Unattached bacteria were removed by thorough washing with PBS, and cells were subjected to immunofluorescence labeling and confocal microscopy as described below.

### Induction and characterization of Citrobacter rodentium infection

All experiments involving *C. rodentium* infection were performed in an ABSL-2 level facility following an established procedure ^41^. Both male and female mice, aged 8–10 weeks, were used for the experiments, with approximately equal distribution of sexes across different experimental groups. Animals were given a streptomycin solution (4 g/L) in drinking water for 24 hours, followed by 24-hour exposure to normal tap water and overnight food deprivation before bacterial gavage. A stationary phase *Citrobacter rodentium* culture was prepared by overnight culture in liquid LB and the concentration of bacterial suspension was determined by measuring its optical density at 600 nm. Animals were gavaged with 200 µL of the bacterial suspension containing 1 x 10^9^ colony-forming units per mouse, or 200 µL of LB medium as a control. The day of the gavage was designated as day 0. Mice were monitored daily for clinical signs of infection, including weight loss, reduced activity, and changes in stool consistency.

To assess bacterial colonization, fecal samples were collected every other day post-infection. Fecal pellets were collected directly into pre-weighed sterile microcentrifuge tubes containing 0.5 ml PBS to allow normalization of bacterial load to the sample mass. Furthermore, after each experiment, approximately 1 cm segments of distal colon and ileum, as well as 0.5 cm segments of cecum tissue, were excised, washed with sterile phosphate-buffered saline (PBS), and placed into pre-weighed sterile tubes.

Samples were homogenized in sterile PBS using a homogenizer with disposable blades ensuring thorough disruption of the tissue and release of bacteria. Homogenates were subjected to ten-fold serial dilutions in sterile PBS, with aliquots of each dilution plated on MacConkey agar plates in triplicate. Plates were incubated at 37°C for 16–18 hours, after which pink lactose-fermenting colonies, characteristic of *C. rodentium*, were counted. The bacterial colony counts were used to calculate CFUs per gram of feces or intestinal tissue, allowing quantitative assessment of the bacterial burden.

### Measurement of epithelial barrier permeability in vivo

Intestinal permeability in NM IIA cKO mice and their control, flox/flox, littermates was performed as previously described ^22, 42^. Briefly, after 3 h of food deprivation, animals were gavaged with fluorescein isothiocyanate (FITC)-labeled dextran (4 kDa; 80 mg/100g body weight) dissolved in phosphate-buffered saline (PBS). Animals were euthanized 3 h later for blood collection via cardiac puncture, blood plasma was obtained by centrifugation, and FITC fluorescence intensity was measured using a Synergy H1 microplate reader (Agilent Technology, Santa Clara, CA) with excitation and emission wavelengths at 495/525 nm. The measured value of dextran-free serum was subtracted from each measurement. The concentration of fluorescent tracer in blood serum was calculated using SigmaPlot v12.5 software, based on a plotted standard curve prepared via serial dilutions of stock solutions of FITC-dextran in PBS.

### Immunoblotting analysis

Mouse colonic segments were harvested, longitudinally dissected and opened, and washed with ice-cold PBS. Epithelial cells were collected by gently scraping the exposed interior with razor blades, then snap frozen into liquid nitrogen for further analysis. Intestinal epithelial scrapings were lysed and homogenized in RIPA buffer containing a protease inhibitor cocktail, phosphatase inhibitor cocktails 2 and 3 and Pefabloc (all from Millipore-Sigma). The samples were diluted with 2x SDS sample loading buffer and boiled. SDS-polyacrylamide gel electrophoresis was conducted using a standard protocol with an equal amount of total protein loaded per lane (10 or 20 µg), followed by immunoblotting on nitrocellulose membrane. Membranes were blocked and sequentially incubated with primary antibodies (at either 1:500 or 1:1,000 dilution) and secondary horseradish peroxidase-conjugated antibodies (at 1:10,000 dilution). The labeled membranes were exposed to the ECL reagent (Millipore-Sigma) with the chemiluminescent signals captured on X-ray film using an automated film processor. Signal intensities were quantified via densitometry using ImageJ software (National Institutes of Health, Bethesda, MD).

### Quantitative real-time RT-PCR

Total RNA was isolated from whole colonic segments of NM IIA cKO and control animals using a RNeasy mini kit (Qiagen, Germantown, MD), followed by DNase treatment to remove genomic DNA. Total RNA (1 µg) was reverse transcribed using an iScript cDNA synthesis kit (Bio-Rad Laboratories). Quantitative real-time RT-PCR was performed using iTaq Universal SYBR Green Supermix (Bio-Rad Laboratories) in a CFX96 Real-time PCR system (Bio-Rad Laboratories). The primer sequences have been published in our previous study ^22^. The threshold cycle number (Ct) for specific genes of interest and a housekeeping gene was determined based on the amplification curve representing a plot of the fluorescent signal intensity versus the cycle number. The relative expression of each gene was calculated by a comparative Ct method based on the inverse proportionality between Ct and the initial template concentration (2^-ΔΔCt^), as previously described ^43^. This method is based on two-step calculations of ΔCt = Ct_target gene_ - Ct_GAPDH_ and ΔΔCt = ΔCt_e_-ΔCt_c_, where index e refers to the sample from any *C. rodentium* or LB-treated NM IIA cKO, or control mice, and index c refers to the sample from a LB-treated control animal assigned as an internal control.

### Immunofluorescence labeling and confocal microscopy

Cells were fixed with 4% paraformaldehyde for 20 minutes at room temperature and permeabilized using 0.5% Triton X-100 for 5 minutes. After washing with Hank’s balanced salt solution (HBSS+), the monolayers were blocked for 60 minutes at room temperature with 1% bovine serum albumin (BSA) in HBSS+ (blocking buffer). Cells were then incubated for 60 minutes with primary antibodies diluted in blocking buffer, washed, and incubated for 60 minutes with Alexa Fluor–conjugated secondary antibodies, rinsed with blocking buffer, and mounted on slides using ProLong™ Gold Antifade Reagent (ThermoFisher Scientific).

F-actin and nuclei were stained with Alexa Fluor 555 (or Alexa 647)–phalloidin and 4′,6-diamidino-2-phenylindole (DAPI), respectively. Fluorescently labeled monolayers were imaged using a Leica SP8 confocal microscope (Wentzler, Germany). Different Alexa fluorophores were imaged sequentially in frame-interlaced mode to prevent spectral overlap. Image processing was performed using Adobe Photoshop (Adobe Systems, San Jose, CA). Individual bacteria or bacterial colonies were quantified using ImageJ. Particle size thresholds were set to 10 pixels² for individual bacteria in the case of HT-29 cells, and 80 pixels² for bacterial colonies associated with Caco-2 cells. To quantify bacterial attachment, the bacterial count in six different microscopic fields per coverslip was averaged to produce a single data point. Three different coverslips per experimental condition were examined and the data obtained in two or three different experiments were combined and analyzed. Numbers of combined experiments are presented in Figure legends.

### Bacterial colony forming assay

Confluent, differentiated IEC monolayers were infected with EPEC at a multiplicity of infection (MOI) of 5:1. The cells were co-cultured with the bacteria in DMEM without antibiotics for 3 hours and then washed three times with PBS to remove non-adherent bacteria. Next, the cells were lysed on ice using 200 µL per well of 1% Triton X-100 in PBS for 10 minutes. Lysates from each well were transferred to microcentrifuge tubes and vortexed. Serial dilutions of the lysates were then plated onto agar plates, which were incubated overnight at 37°C. The next morning, bacterial colonies were counted, and the total number of attached bacteria was calculated as colony forming units per milliliter of cell lysate. Bacterial attachment to three different cell monolayers was counted for each experimental condition. Data combined from 2-3 independent experiments was analyzed.

### Statistics

Data are given as a mean ± SEM. The statistical significance of the difference between the 2 sets of data was evaluated using the two-tailed unpaired Student’s t-test when data were distributed normally. Differences in body weight loss and disease activity index data were tested for statistical significance using one-way ANOVA (SigmaPlot 12.5 package) with Tukey post-hoc test. Statistical significance was accepted at *p* <0.05.

## Results

### Mice with intestinal epithelium-specific knockout of NM IIA show increased susceptibility

Intestinal epithelial cells are known to express two NM II motors, NM IIA and NM IIC, with NM IIA being crucial for establishing the protective intestinal epithelial barrier and limiting mucosal inflammation ^22, 44^. To examine the roles of this cytoskeletal motor in regulating colonization of intestinal epithelium by A/E pathogens *in vivo*, we infected mice with an intestinal epithelial-specific knockout of NM IIA (NM IIA cKO) with *Citrobacter rodentium*. NM IIA flox littermates (NM IIA +/+) served as controls. In agreement with our previous data ^22^, NM IIA cKO mice displayed a selective loss of NM IIA protein expression in the intestinal epithelium without altered expression of NM IIC motor (Figure 1A). NM IIA cKO and control mice were infected with *C. rodentium* at 1×10^9^ CFU per mouse, and bacterial colonization was determined by measuring viable *C. rodentium* from fecal pellets by a colony forming assay. NM IIA cKO mice demonstrated significantly higher bacterial loads on days 2-14 after *C. rodentium* administration as compared to the control animals (Figure 1B). Despite such higher initial colonization in NM II cKO mice, the rate of bacterial clearance appears to be similar between two mouse strains (Figure 1B). Consistently, two weeks post-infection, the amounts of *C. rodentium* recovered from cecal and colonic tissues were not different between NM IIA cKO and their controls, indicating that both animal strains can efficiently clear the pathogen from the intestine (Figure 1C). Interestingly, NM II cKO mice demonstrated exaggerated pathophysiologic responses to the bacterial infection. This included more prominent body weight loss at the early phase of the infection (Figure 1D) and more pronounced colonic crypt hyperplasia after two weeks of *C. rodentium* infection (Figure 1E,F).

**Figure 1.**
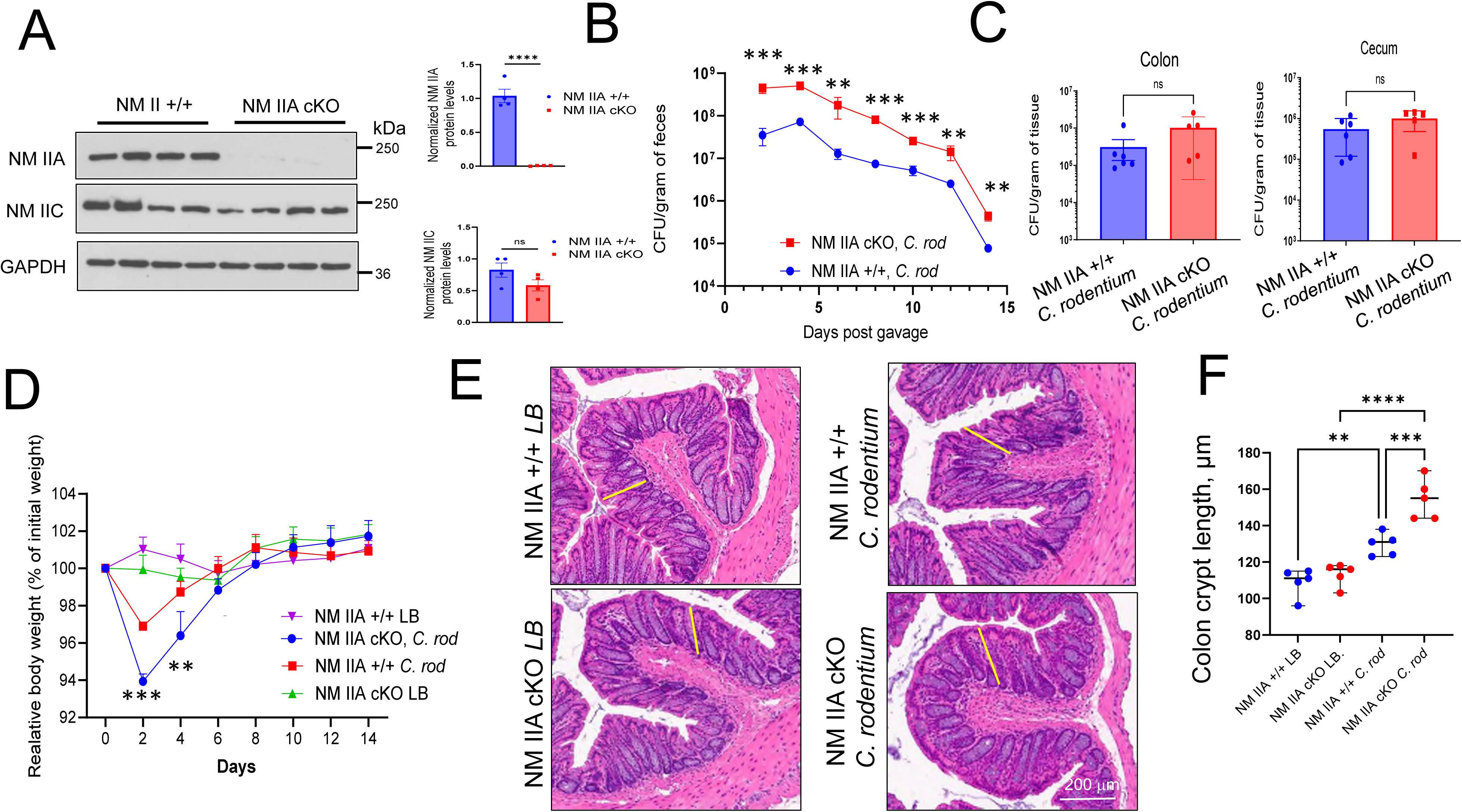
Loss of the intestinal epithelial NM IIA accelerates *Citrobacter rodentium* infection in mice. **(A)** Immunoblotting analysis of NM IIA and NM IIC expression in colonic epithelial scrapes of NM IIA cKO and flox control (+/+) mice. (**B-F**) Control and NM IIA cKO mice were gavaged with *C. rodentium* at 1×10^9 CFU per mouse. (**B**) Bacterial shedding in fecal pellets was examined at the indicated times of the infection. (**C**) Bacterial colonization of the colon and cecum was determined on day 14 of the infection. (**D**) The change in body weight of control and *C. rodentium*-infected animals was determined at different times after bacterial administration. (**E,F**) H&E staining of colonic sections and colonic crypt length was measured in control and *C. rodentium*-infected mice on day 14 after bacterial administration. Mean±SEM, (n=5-6); **p<0.01 *** p<0.001, ****p<0.0001 scale bar=200 μm

Next, we sought to characterize *C. rodentium*-induced intestinal inflammation early post-infection (8 days). In this shorter infection experiment, we observed an approximately 1 log increase in bacterial load in both fecal samples (Supplementary Figure 1) as well as in the colonic and cecal tissues of NM IIA cKO mice (Figure 2A), indicating that loss of NM IIA enhances intestinal colonization by the pathogen. Since A/E bacteria are known to induce breakdown of the intestinal epithelial barrier ^31, 32^, we examined gut barrier integrity in control and NM IIA cKO mice with and without *C. rodentium* infection. Consistent with our previously published data ^22^, NM IIA cKO mice demonstrated increased intestinal permeability under basal conditions (Figure 2B). Furthermore, *C. rodentium* infection triggered further disruption of the gut barrier in NM IIA cKO mice without affecting barrier integrity in control animals (Figure 2B). Next, we evaluated the pathogen-induced mucosal inflammation by measuring mRNA expression of major proinflammatory mediators in colonic tissue samples of NM IIA cKO and control animals. Even unchallenged NM IIA cKO mice showed significant increases in tissue expression of TNFα, and interleukins (IL)-6 and -12, which may suggest a low level of spontaneous inflammation in their intestine (Figure 2C). *C. rodentium* infection resulted in significantly higher mRNA expression of TNFα, IL-6, IL-12, IL-22, interferon-γ, and keratinocyte factor chemokine in colonic tissues of NM IIA cKO mice as compared to their controls (Figure 2C). Together, our data demonstrate that loss of intestinal epithelial NM IIA increases susceptibility to *C. rodentium* infection and promotes pathogen-induced disruption of the gut barrier, mucosal inflammation, and colonic crypt hyperplasia *in vivo*.

**Figure 2.**
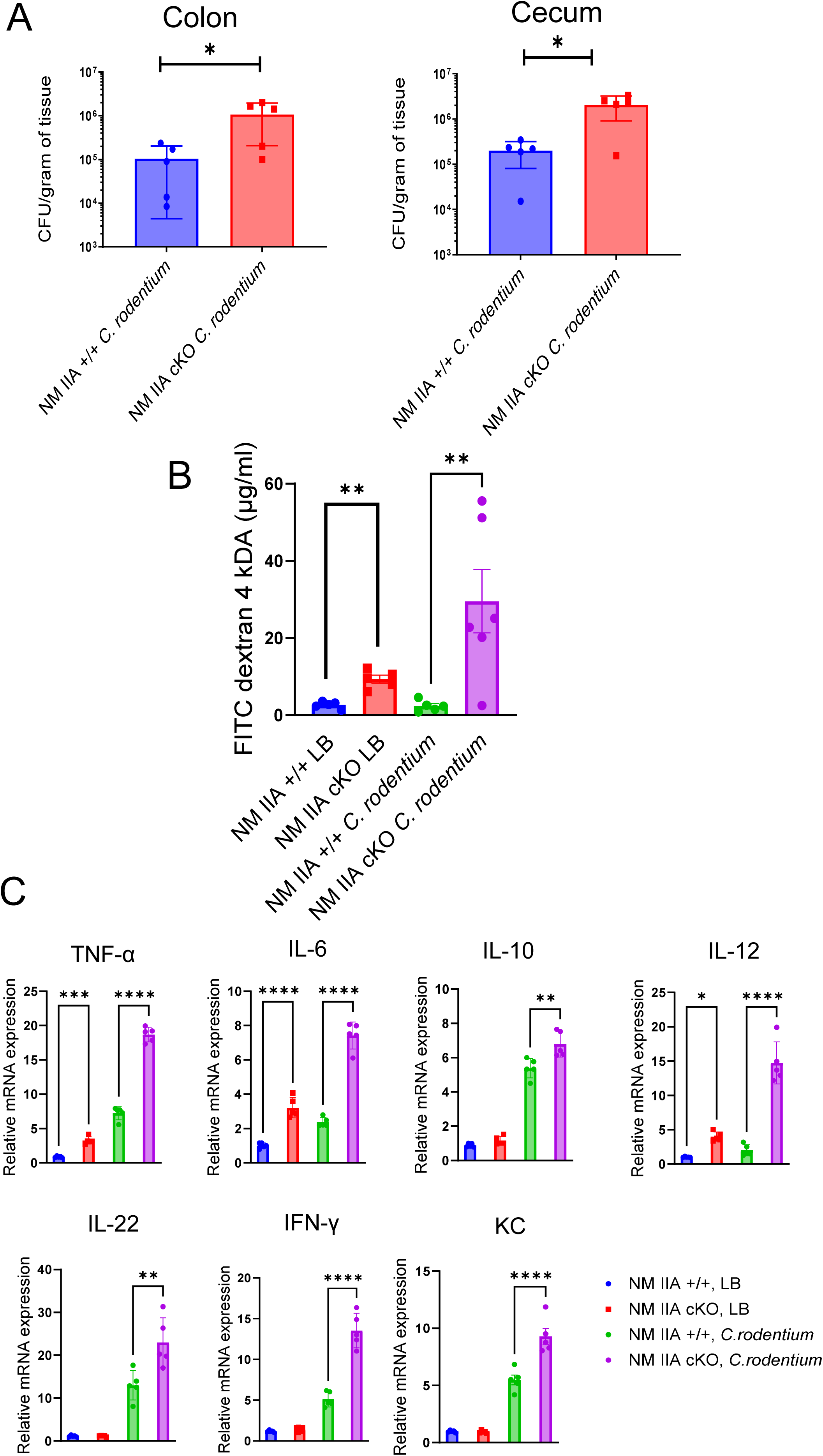
Knockout of intestinal epithelial NM IIA increases animal sensitivity to *C. rodentium*-induced gut barrier disruption and mucosal inflammation. Control and NM IIA cKO mice were gavaged with *C. rodentium* at 1×10^9^ CFU per mouse. (**A**) Bacterial colonization of the colon and cecum was determined on day 8 of the infection. (**B**) The gut-to-blood passage of 4 kDa FITC-dextran was measured in NM IIA cKO and control mice with and without *C. rodentium* infection. Mean±SEM, n=5; **p<0.01. (**C**) mRNA expression of inflammatory markers was measured in colonic tissues of NM IIA cKO and control mice on day 8 of *C. rodentium* infection. Mean±SEM (n = 6); *p < 0.05, **p<0.01 *** p<0.001, ****p<0.0001.

### Pharmacologic or genetic inhibition of NM IIA motor accelerates EPEC attachment to intestinal epithelial cell monolayers in vitro

The increased sensitivity of NM IIA cKO mice to *C. rodentium* infection could be due to enhanced attachment of the A/E pathogen to NM IIA-depleted intestinal epithelium or priming effects of gut barrier disruption and the low-scale mucosal inflammation observed in NM IIA cKO mice (Figure 2). To distinguish between these two possibilities, we adopted a reductionistic *in vitro* approach to examine the attachment of a key human A/E bacterium, EPEC, to HT-29cf8 and Caco-2BBE human colonic epithelial cells. Interactions of EPEC with confluent, differentiated IEC monolayers were examined by either visualizing attached bacteria with immunofluorescence labeling for LPS and confocal microscopy or quantifying live bacteria attached to monolayers by colony-forming assay. NM II motor activity was blocked by a specific pan-NM II inhibitor, blebbistatin ^45^. This was complemented by a gain-of-function approach, utilizing 4-Hydroxyacetophenone (4-HAP) that is known to selectively activate NM IIB and NM IIC activity without affecting NM IIA ^46^. Blebbistatin treatment significantly increased EPEC attachment to HT-29 (Figure 3A, B) and Caco-2 (Figure 3C, D) cells in both adhesion assays. By contrast, 4-HAP treatment did not significantly change EPEC interactions with IECs.

**Figure 3.**
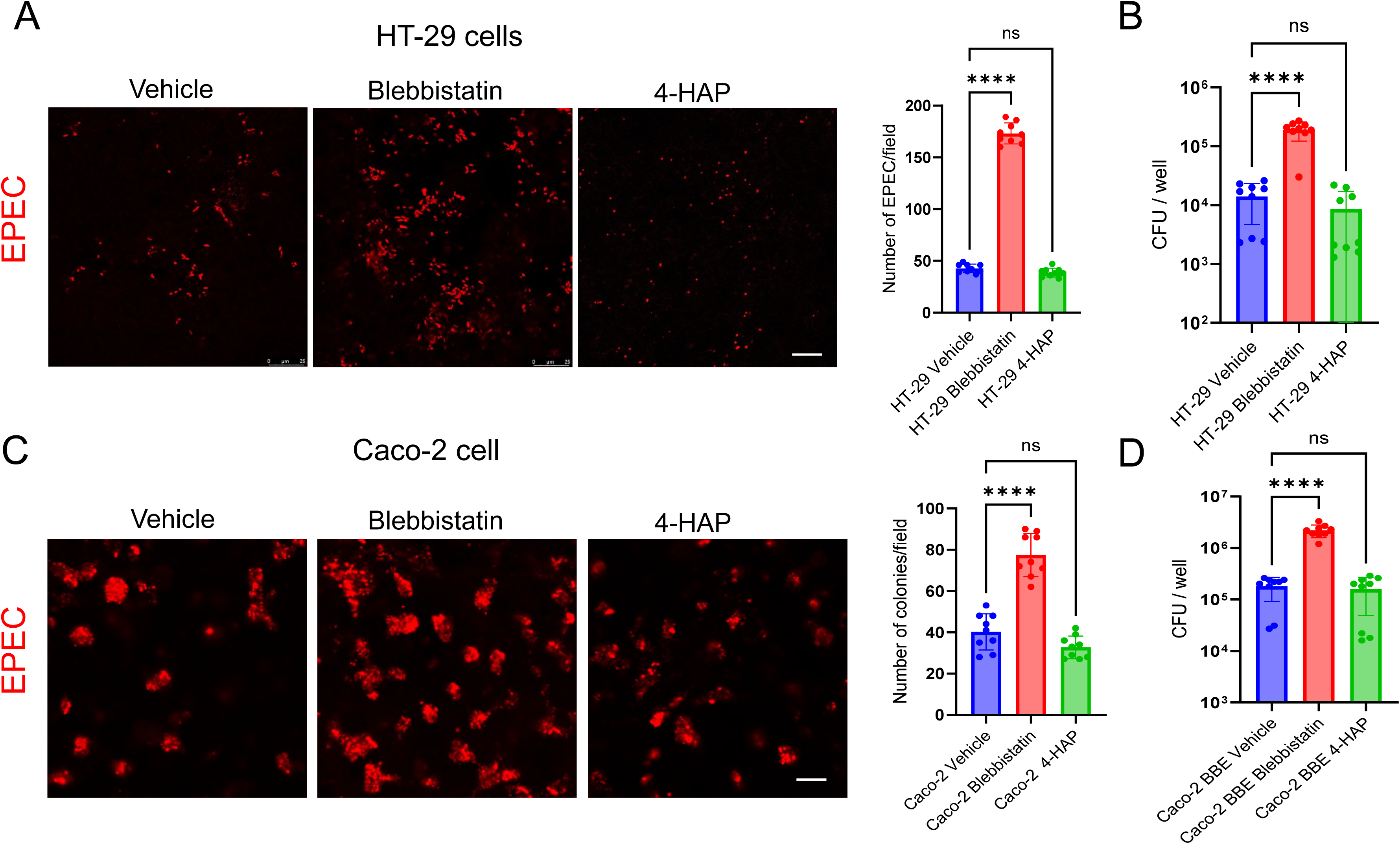
Pharmacological inhibition of NM II motor activity promotes EPEC attachment to model intestinal epithelial cell monolayers *in vitro*. Confluent differentiated HT-29cf8 (**A,B**) Caco-2BBE (**C,D**) cell monolayers were exposed to EPEC (MOI 5:1) for 3 h in the presence of either vehicle, blebbistatin (50 µM), or 4-HAP (100 µM). Bacterial attachment to IEC was determined by either immunofluorescence labeling/confocal microscopy of LPS (**A,C**) or colony forming assay (**B,D**). Mean±SEM of combined data from three independent experiments (n=9); **p<0.01, **** p<0.001; scale bar=20 μm.

Next, we examined whether NM IIA is required for restricting EPEC attachment to model IEC monolayers. To examine the functional role of this cytoskeletal motor, we created IEC lines with stable CRISPR/Cas9-induced knockout of NM IIA. Using two different sgRNAs, we achieved a marked reduction (up to 99%) of NM IIA protein expression in HT-29 and Caco-2 cells without affecting NM IIC levels (Figure 4A,D). Such NM IIA knockout significantly upregulated EPEC attachment in both HT-29 (Figure 4B,C) and Caco-2 cell monolayers (Figure 4D,E), thereby recapitulating the results obtained with *C. rodentium* infection of NM IIA cKO mice.

**Figure 4.**
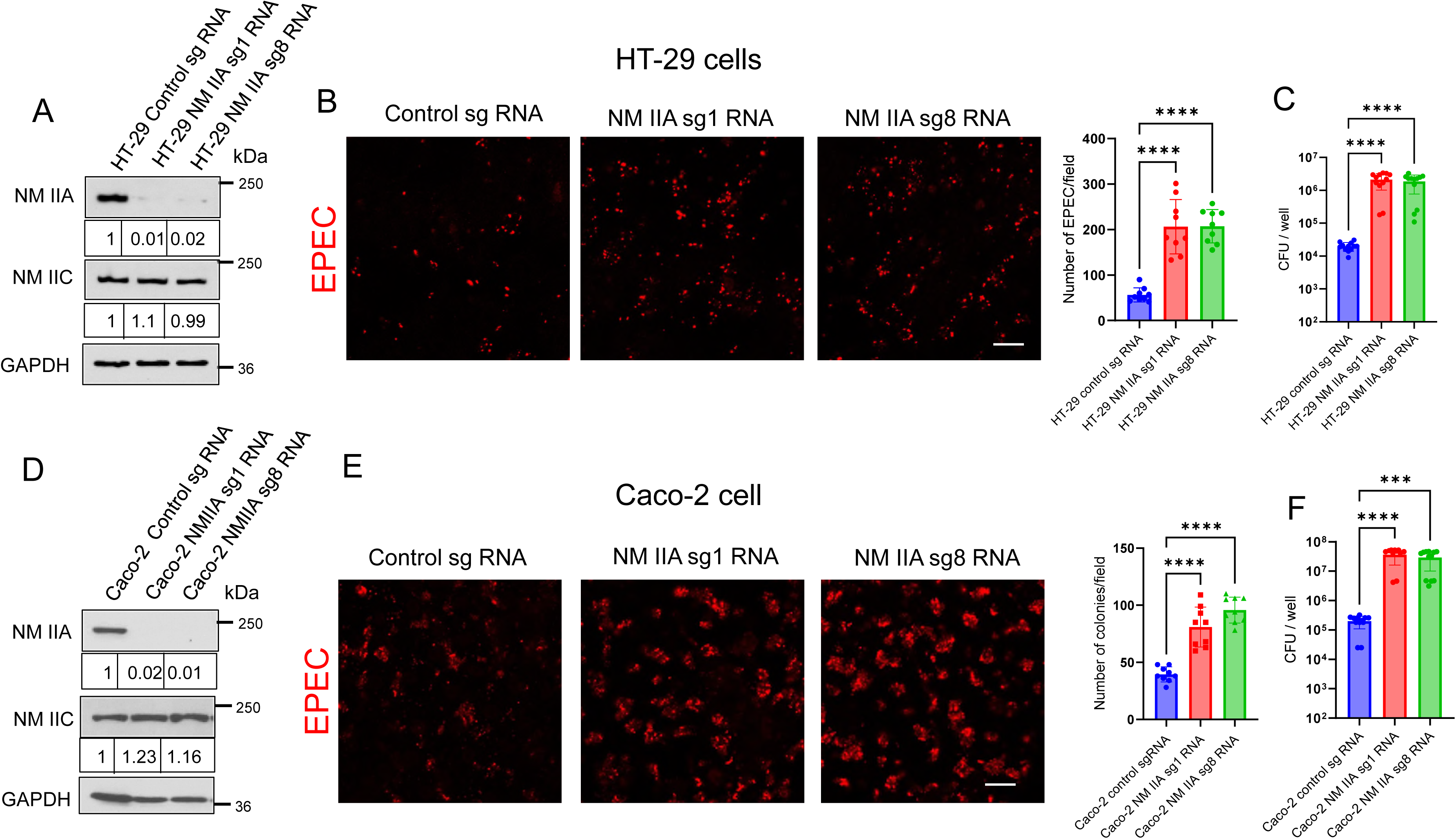
CRISPR-Cas9-mediated knockout of NM IIA promotes EPEC attachment to model intestinal epithelial cell monolayers *in vitro*. (**A,D**) Immunoblotting analysis of NM IIA and NM IIC expression in HT-29cf8 and Caco-2BBE cells with CRISPR/Cas9-mediated knockout of NM IIA using two different sgRNAs. (**B-E**). Control and NM IIA knockout HT-29 (**B,C**) and Caco-2 (**E,F**) cells were infected with EPEC (MOI 5) for 3 h. Bacterial attachment to IEC was determined by either immunofluorescence labeling/confocal microscopy of LPS (**B,E**) or colony forming assay (**C,F**). Mean±SEM of combined data from three independent experiments (n=9); **p<0.01, **** p<0.001; scale bar=20 μm.

### Loss of NM IIC expression does not affect A/E bacterial interactions with intestinal epithelial cells in vitro and in vivo

Next, we sought to investigate if another major IEC actin motor, NM IIC, plays role in mediating A/E pathogens interactions with intestinal epithelium. CRISPR/Cas9-mediated knockout of NM IIC caused a significant decrease in its protein level in Caco-2 cells, while NM IIA expression remained unaffected (Figure 5A). However, unlike NM IIA knockout, loss of NM IIC did not impact EPEC attachment to IEC monolayers as shown by immunofluorescence labeling/confocal microscopy of attached bacteria (Figure 5B) or bacterial colony forming assay (Figure 5C). We sought to determine the *in vivo* relevance of this finding by examining *C. rodentium* infection in mice with total knockout of NM IIC (NM IIC tKO). Predictably, NM IIC tKO mice showed complete loss of NM IIC protein expression in the colonic epithelium, while NM IIA levels remained unchanged (Figure 5D). Both NM IIC tKO and control mice demonstrated similar responses to *C. rodentium* infection, based on their similar bacterial loads in feces and intestinal tissues (Figure 5E,F), no significant body weight loss (Figure 5G), and similar levels of colonic crypt hyperplasia (Figure 5H). Together, these data suggest that the NM IIC motor does not regulate interactions of A/E pathogens with the intestinal mucosa *in vivo* and *in vitro*.

**Figure 5.**
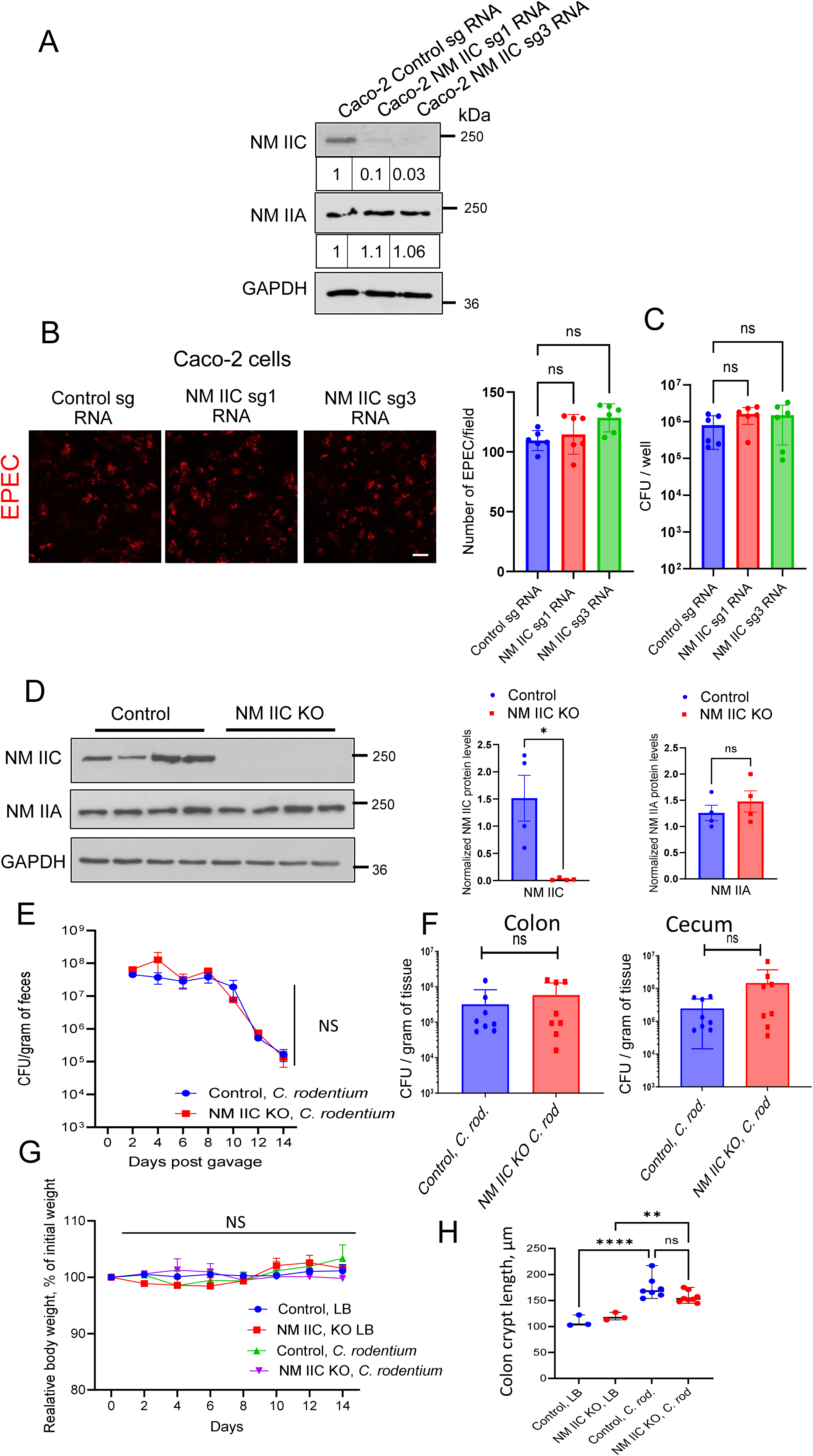
Loss of intestinal epithelial NM IIC does not affect A/E bacterial interactions with intestinal epithelium *in vitro* and *in vivo*. **(A)** Immunoblotting analysis of NM IIC and NM IIA expression in Caco-2BBE cells with CRISPR/Cas9-mediated knockout of NM IIC using two different sgRNAs. (**B,C**). Control and NM IIC knockout Caco-2 cells were infected with EPEC (MOI 5:1) for 3 h. Bacterial attachment to IEC was determined by either immunofluorescence labeling/confocal microscopy of LPS (**B**) or colony forming assay (**C**). Mean±SEM of combined data from two independent experiments (n=6). (**D**) Immunoblotting analysis of NM IIC and NM IIA expression in colonic epithelial scrapes of NM IIC knockout and control mice. (**E-H**) Control and NM IIC KO mice were gavaged with *C. rodentium* at 1×10^9^ CFU per mouse. (**E**) Bacterial shedding in fecal pellets was examined at the indicated times of the infection (**F**) Bacterial colonization of the colon and cecum was determined on day 14 of the infection (**G**) The change in body weight of control and *C. rodentium*-infected animals determined over time. (**H**) Colonic crypt length was measured in control and *C. rodentium*-infected mice on day 14 after bacterial administration. Mean±SEM, n=3-7; **p<0.01, **** p<0.001; scale bar=20 μm

### Motor domain activity is essential for NM IIA-dependent regulation of A/E bacteria interactions with the intestinal epithelium

Given the observed unique roles of NM IIA in limiting A/E bacterial attachment to IECs, we next sought to understand what properties of this cytoskeletal protein are essential for controlling host-bacterial interactions. Like other conventional myosins, NM IIA has dual functionality as an actin motor and cross-linking protein ^19, 20^. Its motor function is determined by the N-terminal globular domain that interacts with actin filaments and possesses ATPase activity ^19, 20^. Several point mutations in the N-terminal domain were shown to inhibit the motor activity of NM II ^47, 48^. We examined whether the motor activity of NM IIA is essential for controlling A/E bacterial interactions with mammalian cells by utilizing two motor domain mutations, R702C and N93K. The GFP-tagged mutants, along with wildtype NM IIA, were transiently expressed in Cos-7 epithelial cells that lack endogenous NM IIA. EPEC attachment to GFP-NM IIA expressing cells was determined by immunolabeling and confocal microscopy. Bacterial attachment to Cos-7 cells expressing NM IIA mutants was significantly higher than attachment to WT NM IIA-expressing cells (Figure 6A). To establish the *in vivo* relevance of this finding, we used transgenic mouse strains expressing GFP-labeled WT human NM IIA or its R702C mutant under control of the endogenous mouse promoter ^34^. GFP-NM IIA WT and GFP-NM IIA R702C mice were infected with *C. rodentium* for 14 days, and the bacterial infection was characterized as described above. Compared to WT controls, the NM IIA R702C mice demonstrated a 1 log higher fecal bacterial loads during the infection (Figure 6B) and significantly elevated *C. rodentium* levels in cecal tissue 14 days after bacterial administration (Figure 6C). Furthermore, the mutant mice also displayed more pronounced body weight loss (Figure 6D) and colonic crypt hyperplasia compared to their WT controls (Figure 6E,F), thereby recapitulating higher sensitivity to *C. rodentium* infection observed in the NM IIA cKO mice. Collectively, these results indicate that the motor activity of NM IIA is essential for limiting A/E bacterial interactions with the intestinal epithelium *in vivo* and *in vitro*.

**Figure 6.**
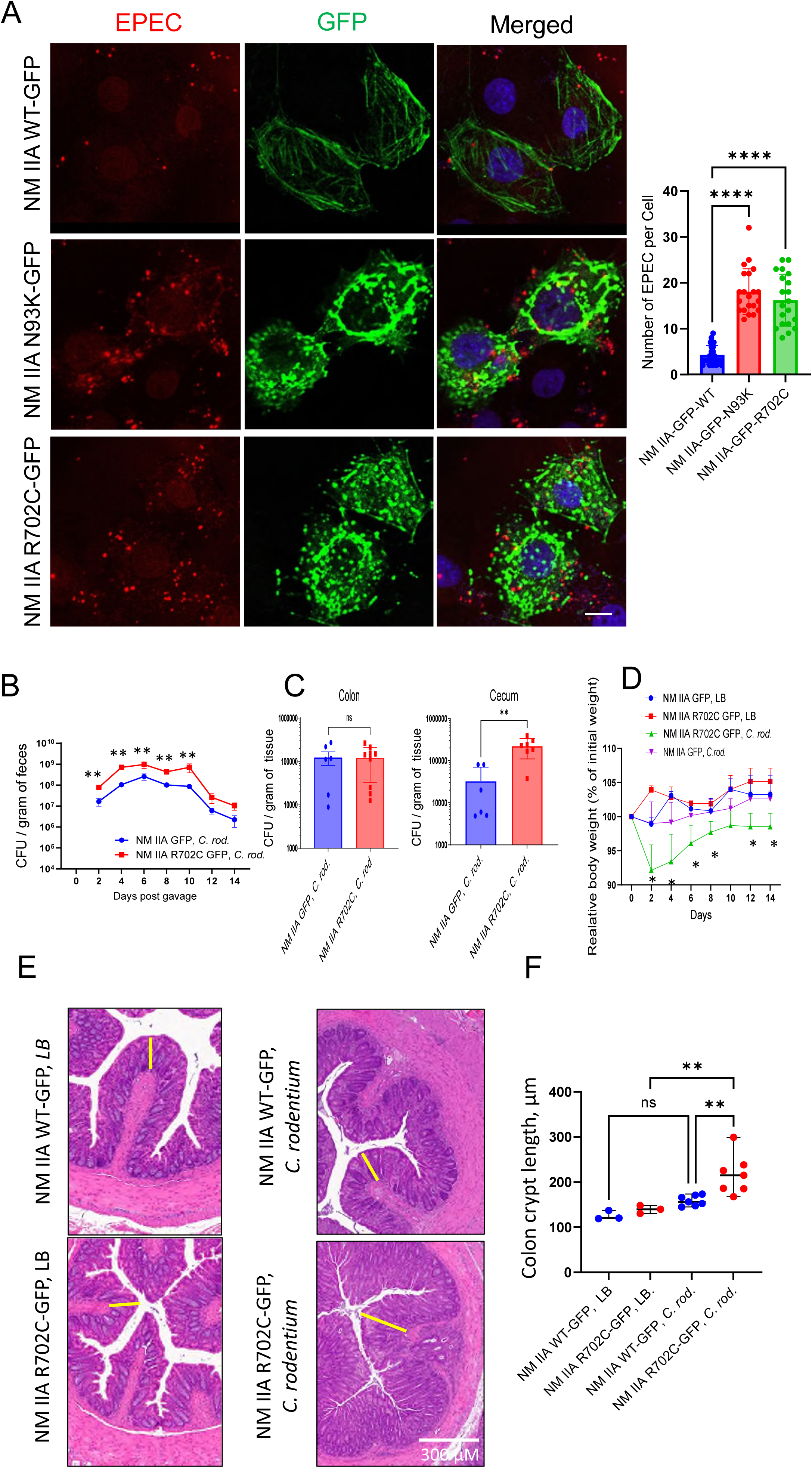
Mutations in the motor domain of NM IIA promote A/E bacterial interactions with intestinal epithelium *in vitro* and *in vivo*. **(A)** EPEC attachment to Cos-7 cells (MOI 5:1 for 3 h) transfected with GFP labeled WT NM IIA or NM IIA R702C and N93K mutants was determined by immunofluorescence labeling and confocal microscopy Mean±SEM, n=20-25 individual cells per group. Data is representative of 3 independent experiments. (**B-F**) Transgenic mice expressing either WT GFP-NM IIA or GFP-NM IIA R702C mutant were gavaged with *C. rodentium* at 1×10^9^ CFU per mouse. (**B**) Bacterial shedding in fecal pellets was examined at the indicated times of the infection. (**C**) Bacterial colonization of the colon and cecum was determined on day 14 of the infection. (**D**) The body weight of control and *C. rodentium*-infected animals was measured at different times after bacterial administration. (**E,F**) Colonic crypt length was measured in control and *C. rodentium*-infected mice on day 14 after bacterial administration. Mean±SEM, n=3-7; *p<0.05, **p<0.01, **** p<0.001; scale bar=10 μm

### NM IIA controls Tir-dependent EPEC attachment to IEC monolayers

Next, we sought to investigate the mechanisms by which NM IIA motor attenuates the attachment of A/E bacteria to the intestinal epithelium. We rationalized that NM II most likely regulates the actin cytoskeletal remodeling induced by bacterial attachment, more specifically, assembly of F-actin pedestals. Indeed, fluorescence labeling and confocal microscopy revealed the formation of significantly larger F-actin pedestals at the EPEC attachment sites in NM IIA-deficient IEC monolayers compared to control cells (Figure 7A). To decisively prove that NM IIA controls EPEC attachment to IECs in an actin pedestal assembly-dependent fashion, we investigated cellular interactions with EPEC mutants that lack a key actin pedestal-forming effector, Tir ^49^. Specifically, we compared cell attachment of the bacterial strain with total Tir deletion (EPEC-Δ*tir*) and EPEC strains reconstituted with either wild-type Tir (EPEC-TirWT) or with mutations of two key residues (Y454F/Y474F) responsible for triggering actin polymerization and pedestal assembly (EPEC-TirY454F/Y474F) ^35–38^. As an additional control, we used an EPEC mutant with a deleted Type III secretion system that is unable to introduce any bacterial effectors into the host cells ^37, 38^. As expected, the infection of HT-29 cells monolayers with the EPEC-TirWT strain resulted in robust assembly of F-actin pedestals beneath the bacterial colonies (Supplementary Figure 2, arrows). By contrast, IEC infection with either the EPEC-Δ*tir* or the EPEC-TirY454F/Y474F mutants did not result in pedestal assembly (Supplementary Figure 2, arrowheads). NM IIA knockout IEC monolayers demonstrated a significantly increased attachment of EPEC-TirWT (Figure 7B,C), recapitulating our results obtained with the parental EPEC strain (Figure 4). However, IEC attachment of both pedestal formation-deficient Tir mutants was markedly lower when compared to the EPEC-TirWT (Figure 7B,C). More importantly, NM IIA knockout did not enhance the cell adhesion of the EPEC-Δ*tir* or the EPEC-TirY454F/Y474F mutants (Figure 7B,C). Similarly, the T3SS-deficient EPEC mutant showed very low IEC attachment that was not affected by NM IIA knockout (Supplementary Figure 3). These data indicate that NM IIA limits EPEC attachment to IECs by attenuating bacterial-induced actin pedestal assembly.

**Figure 7.**
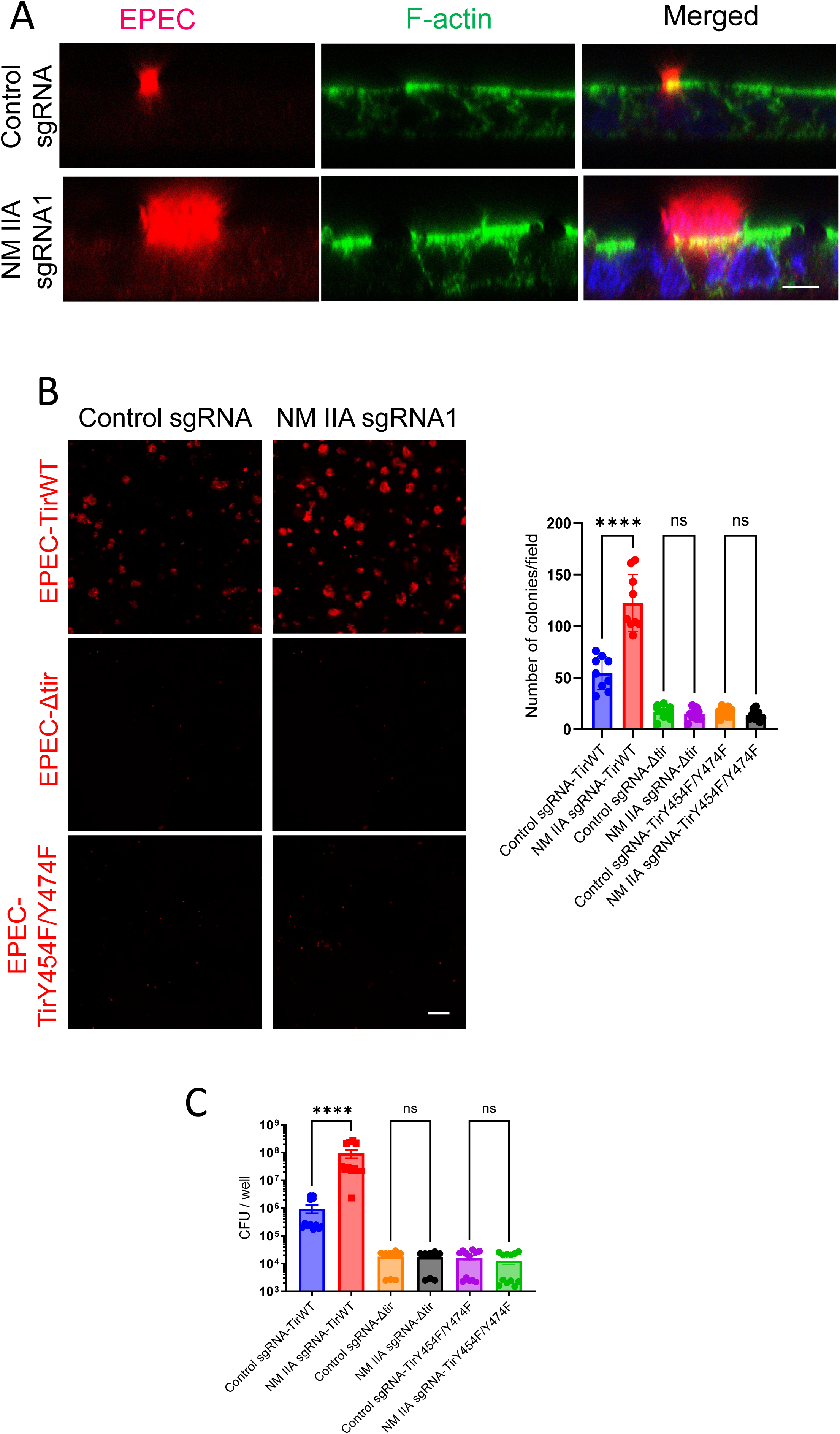
NM IIA does not affect IEC attachment of EPEC pedestal formation deficient mutants. **(A)** Control and NM IIA knockout Caco2-BBE cells were infected with EPEC (MOI 5:1) for 3h. Cells were fixed and fluorescently labeled for F-actin (green) and bacteria (red). Arrow and arrowhead point at F-actin pedestal under bacterial colonies in control and NM-IIA knockout cells, respectively. Scale bar=10μm. (**B,C**). Control and NM IIA knockout HT-29 cells were infected with a Tir-deficient EPEC strain (EPEC-Δ*tir*) or bacterial strain reconstituted with wild-type Tir (EPEC-TirWT) or its Y454F/Y474F mutant (EPEC-TirY454F/Y474F). IECs were exposed to MOI 5:1 of the bacteria for 3 h. Bacterial attachment to IECs was determined by either immunofluorescence labeling/confocal microscopy of LPS (**B**) or colony forming assay (**C**). Mean±SEM of combined data from three independent experiments (n=9); **** p<0.001; scale bar=20μm.

### Loss of NM IIA inhibits EPEC-induced assembly of basal stress fibers in intestinal epithelial cells

How could NM IIA diminish the actin pedestal assembly of EPEC-infected IECs? Immunofluorescence labeling and confocal microscopy did not detect specific accumulation of NM IIA in the apical pedestals (Supplementary Figure 4). Paradoxically, NM IIC was found to be enriched in these structures (Supplementary Figure 4, arrow). Lack of specific pedestal accumulation of NM IIA indicates this motor is unlikely to exert local control of the actin filament structure and dynamics in the pedestals. We rationalized that NM IIA may regulate EPEC-induced global remodeling of the actin cytoskeleton, thereby limiting pedestal assembly. In addition to triggering formation of the apical actin pedestals, A/E bacteria are known to induce assembly of basal stress fibers in host cells ^50, 51^. Therefore, one could envision these pathogen-induced apical and basal cytoskeletal structures antagonize each other’s assembly by competing for the monomeric actin, already in short supply even in normal cells ^52^. Since NM IIA is localized at the EPEC-induced basal stress fibers (Supplementary Figure 5), it can promote assembly of these actin structures and decrease the availability of monomeric actin for the apical pedestal formation.

We first tested the monomeric actin supply requirement by treating IECs with two small molecular compounds, known to regulate the equilibrium between monomeric and polymeric actin by different mechanisms. Latrunculin A decreases the supply of monomeric actin by sequestering such monomers and preventing their incorporation into the actin filaments ^53, 54^. Cytochalasin D increases the supply of monomeric actin by promoting filament depolymerization at the pointed end, while still allowing short filament assembly ^55, 56^. While Latrunculin A treatment significantly reduced EPEC attachment to Caco-2 cell monolayers, exposure to cytochalasin D increased bacterial adhesion to IEC (Supplementary Figure 6). This indicates that bacterial interactions with IEC can be regulated by the availability of monomeric actin. Finally, we examined the formation of stress fibers in EPEC-exposed control and NM IIA-depleted IEC. Consistent with the data obtained in other experimental systems ^50, 51^, EPEC infection of control HT-29 and Caco-2 cells induced a robust assembly of basal actin filament bundles (Figure 8, arrows). By contrast, a little basal F-actin bundle assembly was detected in EPEC-exposed NM IIA knockout IEC (Figure 8, arrowheads). Together, these data suggest that loss of NM IIA could redirect the pathogen-induced actin cytoskeletal assembly from the basal stress fibers toward the apical pedestals, thereby promoting bacterial attachment to IEC.

**Figure 8.**
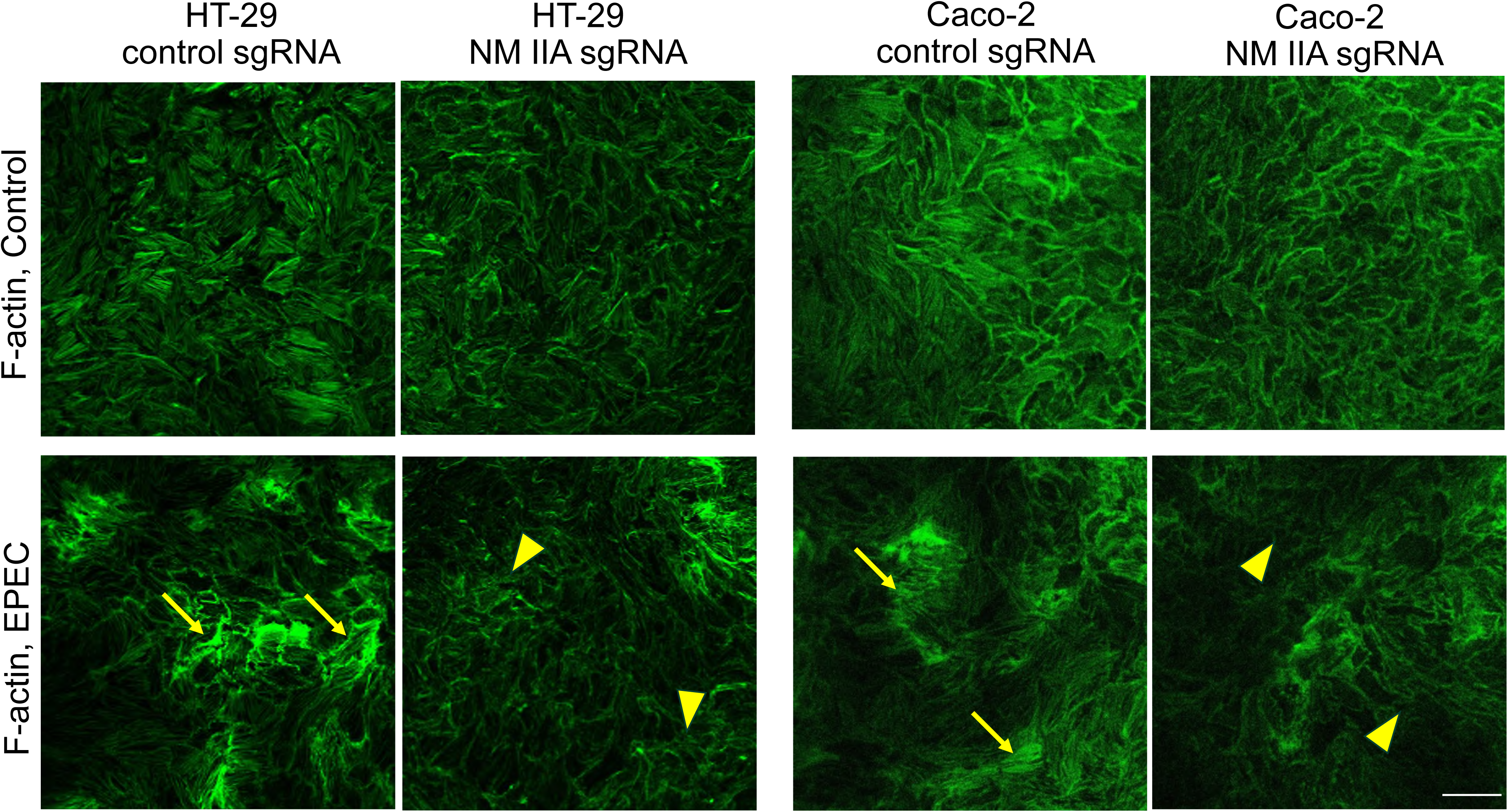
Loss of NM IIA attenuated basal stress fiber assembly in EPEC-infected intestinal epithelial cells. Control and NM IIA knockout HT-29cf8 (**A**) and Caco-2BBE (**B**) cells were infected with EPEC (MOI 5:1) for 3h. Cells were fixed and fluorescently labeled for F-actin (green). Arrows point at basal stress fiber assembly in EPEC-infected control epithelial cell monolayers. Arrowheads indicate poor stress fiber assembly in the infected NM IIA knockout cells. Representative images and quantitation of F-actin signal intensities are shown. Mean±SEM, n=4; **p<0.01, **** p<0.001; scale bar=20 μm.

## Discussion

Attaching-effacing bacteria colonize mammalian host cells by triggering massive remodeling of the actin cytoskeleton, resulting in the assembly of actin pedestals beneath bacterial attachment sites ^2, 10, 15^. Formation of such apical pedestals involves a large number of different actin-binding and regulating proteins; however, functional roles of only a small fraction of these proteins in regulating host-pathogen interactions have been defined ^16, 17^. Our study demonstrates for the first time that a key F-actin motor, NM IIA, acts as a negative regulator of A/E bacterial colonization of intestinal epithelial cells by interfering with actin pedestal assembly.

Our conclusion, highlighting NM IIA as a negative regulator of A/E pathogen interactions with intestinal epithelial cells, is based on the observation that either pharmacologic or genetic inhibition of this cytoskeletal motor increased EPEC attachment to model IEC monolayers *in vitro* (Figures 3 & 4). The physiological significance of this regulatory function of NM II was supported by our data showing the increased *C. rodentium* colonization in mice with intestinal epithelium-specific deletion of NM IIA (Figure 1). Specifically, our findings implicate the NM IIA paralog in regulating A/E pathogen interactions with intestinal epithelial cells. Indeed, genetic knockout of NM IIA in model IEC or mouse intestinal mucosa promoted *C. rodentium* and EPEC infections (Figures 1 & 4). Expression of the NM IIA R702C mutant with diminished motor activity recapitulated the effects of NM IIA deletions in enhancing A/E bacterial interactions with intestinal epithelium (Figure 6), further supporting the unique roles of NM IIA motor in controlling enteric bacterial infection.

In contrast to the roles of NM IIA, we did not find NM IIC involvement in A/E bacterial infection, which is unexpected since this protein is ideally positioned to control enteric pathogen interactions with intestinal epithelium. It selectively localizes at the apical pole of polarized IECs, being enriched in the perijunctional actomyosin belt ^22, 57^. Furthermore, NM IIC is known to regulate microvilli dynamics at the epithelial surface ^23^ and may therefore control microvilli effacement that is induced by the attaching pathogens. Finally, our data suggests that NM IIC is enriched at EPEC pedestals (Supplementary Figure 4). Yet, either loss of NM IIC expression or its pharmacologic activation had no effects on EPEC-interaction with IEC monolayers *in vitro* (Figure 3 and Figure 5B,C), whereas total knockout of NM IIC in mice did not affect animal responses to *C. rodentium* colonization (Figure 5E-H).

Our study could help to resolve a long-standing uncertainty regarding the functional roles of NM II during A/E bacterial infections, which likely reflects the use of suboptimal experimental tools to interfere with myosin activity. Thus, previous pharmacological inhibition of MLC phosphorylation did not affect actin filament dynamics within the EPEC pedestals or pedestal motility on the epithelial surfaces ^58^. By contrast, studies utilizing butanedione monoxime (BDM) to block the ATPase activity of myosin II reported substantial elongation of actin pedestals in EPEC and EHEC-infected epithelial cells ^24, 58^. However, BDM is not a specific NM II inhibitor since it could also disrupt the actin filament turnover ^59^. Therefore, the pedestal elongating activity of BDM could involve myosin-independent mechanisms. Subsequently, a report identified an EPEC effector, EspB, as an NM II-binding protein ^26^. Deletion of EspB was found to inhibit EPEC-induced microvilli effacement in Caco-2 cells, and attenuate intestinal colonization by *C. rodentium* in mice ^26^. However, in addition to NM II, EspB also interacts with actin^60^ and the unconventional myosins 1c, 5, 6, and 10 ^26^. Therefore, the decreased host colonization by the EPEC mutant with deleted EspB cannot be attributed to the selective inhibition of NM II activity. Our study that combines pharmacologic and genetic inhibition of NM II motors demonstrates the increased intestinal epithelial colonization by A/E pathogens when NM IIA activity is compromised, both in cultured IEC *in vitro* and mouse intestinal mucosa *in vivo*.

Importantly, our data suggests that NM IIA-dependent regulation of EPEC attachment to model IEC monolayers involves the assembly of bacterial pedestals. This conclusion is based on the observation of larger pedestals induced by EPEC in NM IIA-deficient epithelial cells (Figure 7A). Furthermore, NM IIA deletion promoted attachment of only EPEC strains with the functional Tir effector that is capable of initiating actin pedestal assembly. By contrast, NM IIA inhibition did not affect the attachment of EPEC strains with either deletion of Tir or expressing the pedestal formation-deficient Tir mutant (Figure 7B,C).

How could NM IIA limit assembly of apical actin pedestals? We did not find selective recruitment of NM IIA to the pedestals in IECs (Supplementary Figure 4), indicating that this actin motor affects pedestal formation remotely. Such remote actions are likely to be linked to controlling the remodeling of other parts of the epithelial actin cytoskeleton triggered by attaching pathogens. Specifically, our data suggests that NM IIA could be essential to pathogen-induced assembly of basal stress fibers. The stress fiber assembly may aid bacterial colonization by increasing IEC attachment to the extracellular matrix and slowing down the extrusion of infected cells from epithelial monolayers. However, the simultaneous assembly of the apical pedestals and basal stress fibers could be antagonistic due to competition for monomeric actin, which is required to build both structures. Indeed, sequestration of monomeric actin with Latrunculin A inhibited, while an increase in monomeric actin supply by Cytochalasin D treatment accelerated EPEC attachment to IEC monolayers (Supplementary Figure 6). The increased bacterial attachment in Cytochalasin-treated cells could be considered paradoxical since this compound is believed to prevent actin polymerization by capping the fast-growing barbed ends of actin filaments. However, a recent study that revisited the cellular activity of cytochalasins revealed that they do not completely block but rather remodel actin filament growth and even increase their association with actin elongating Ena/VASP proteins ^55^. The combined effects of the enhanced actin monomer supply and the increased recruitment of Ena/VASP proteins are likely to explain the increased EPEC attachment to Cytochalasin D-treated IECs.

Robust inductions of basal actin stress fibers were evident in EPEC-infected control HT-29 cells (Figure 8) and these stress fibers were enriched in NM IIA (Supplementary Figure 5). Importantly, assembly of EPEC-induced stress fibers was attenuated in NM IIA-deficient IEC monolayers (Figure 8). Taken together, these data suggest that NM IIA is essential for the assembly/stabilization of basal stress fibers induced by A/E pathogens. We hypothesize that attenuated stress fiber assembly in NM IIA-depleted IECs increases the availability of monomeric actin for the formation of apical actin pedestals, leading to the increased A/E pathogen attachment. This idea is supported by a previous study showing that an EPEC effector protein, EspM, that induces stress fiber assembly, blocks actin pedestal formation in the infected mammalian cells ^61^.

In conclusion, the present study revealed a novel role of NM IIA motor in inhibiting interactions of A/E pathogens with intestinal epithelium in vitro and in vivo. NM IIA serves as an essential regulator of bacteria-induced global remodeling of the actin cytoskeleton in IECs by attenuating assembly of apical actin pedestals and accelerating formation of basal stress fibers. This function requires the motor activity of NM IIA and is not shared by its close paralog, NM IIC. Our findings open an opportunity for using pharmacological activation of NM II motors in order to decrease intestinal colonization by A/E pathogens.

## Acknowledgments

This work was supported by National Institutes of Health grant R01DK131550 to A.I.I. Confocal microscopy performed at the Cleveland Clinic Research Digital Imaging Microscopy Core utilized the Leica SP8 confocal microscope that was purchased with funding from the National Institutes of Health SIG grant 1S10OD019972-01. We thank Dr. Michelle Dziejman (University of Rochester School of Medicine) for the insightful comments on the manuscript.

## Author Contributions

AII conceptualized the study, acquired funding, designed the experiments, interpreted the data, wrote the manuscript, and supervised the study. NGN performed most of the experiments, analyzed and interpreted the data, and contributed to the writing of the manuscript, AZ and SL managed mice colonies and contributed to all animal studies. AMM performed western blot experiments and densitometric data analysis. CM designed animal studies and edited the manuscript. KC generated bacterial mutants, contributed to the study design and manuscript writing. JAH and VMF provided mouse strain and edited the manuscript.

## Data availability

Data available withing the article and its supplementary materials. All relevant original data is available on request from the corresponding author.

## List of abbreviations

A/E: attaching/effacing
cKO: conditional knockout
EPEC: Enteropathogenic *Escherichia coli*
IECs: intestinal epithelial cells
NM II: non-muscle myosin II
T3SS: Type III secretion system
tKO: total knockout

