## Supplementary figures for "Myosin IIA motor regulates attaching-effacing bacteria interactions with intestinal epithelium"

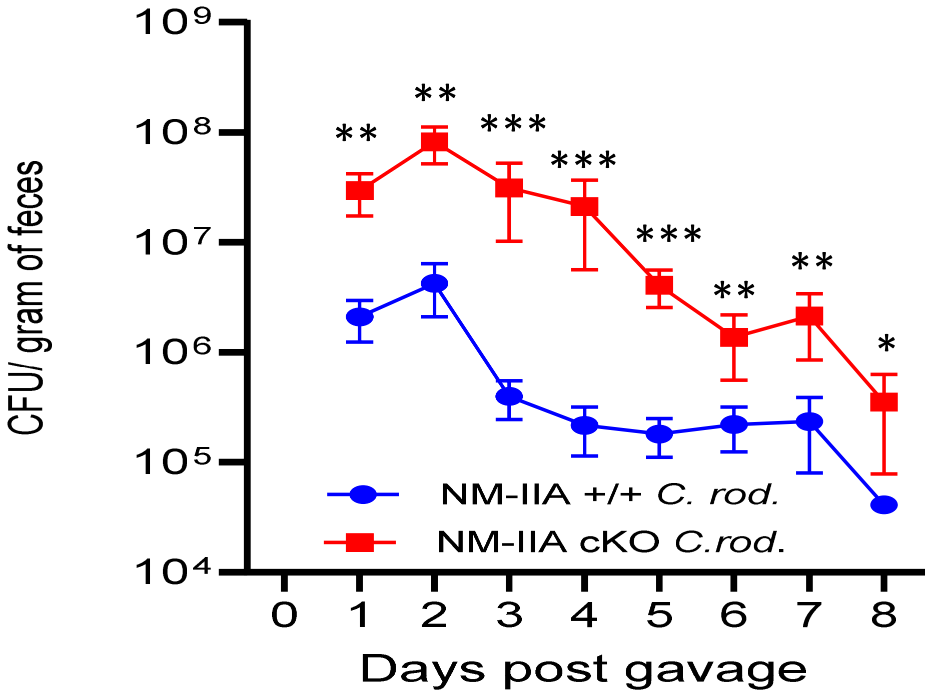


**Supplementary Figure 1. Increased *Citrobacter rodentium* colonization of NM IIA cKO mice during a short-term infection.**

Control and NM IIA cKO mice were gavaged with *C. rodentium* at 1x10^9 CFU per mouse. Bacterial shedding in fecal pellets was examined during 8 days of the infection. Mean±SEM, n=5; **p<0.01, ***p<0.001.

Supplementary. Figure 1

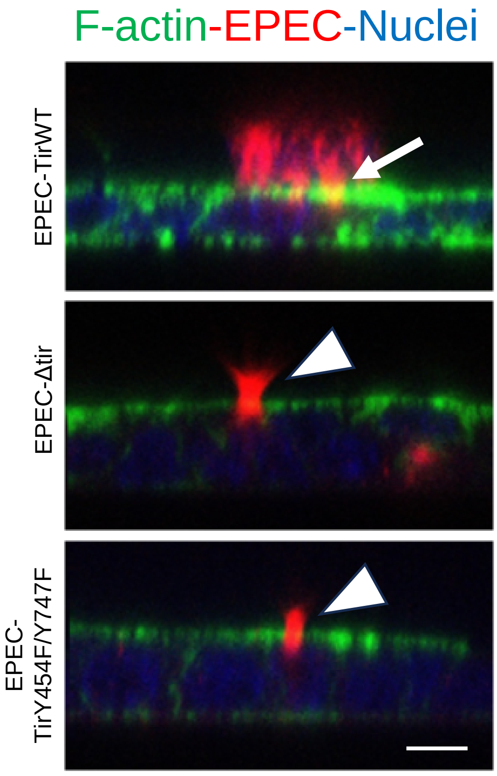


**Supplemental Figure 2: Tir-deficient and Tir phosphorylation-deficient EPEC mutants do not form actin pedestals after infection of IEC monolayers.**

Confluent differentiated HT-29 cell monolayers were infected at (MOI 5:1 for 3 h) with the Tir-deficient EPEC strain (EPEC-Δtir) or the bacterial strain reconstituted with wild-type Tir (EPEC-TirWT) and its Y/Y mutant (EPEC-TirY454F/Y747F). Cells were fixed and fluorescently labeled for F-actin (green) and bacteria (red). Arrows point at F-actin pedestal under bacterial colonies in EPEC WT Tir-infected cells. Arrowheads point at a lack of pedestal assembly in epithelial cells infected with two Tir dysfunctional mutants. Scale bar=10 μM.

Supplementary Figure 2

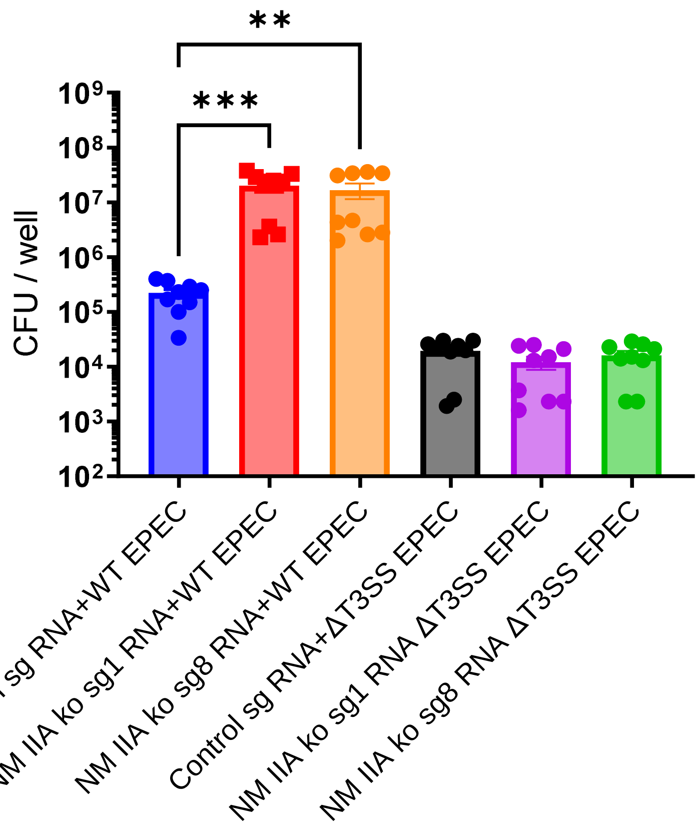


**Supplemental Figure 3: Low, NM IIA-independent IEC adhesion of the EPEC mutant lacking the Type III secretion system.**
Control or NM IIA-deficient HT-29 cell monolayers were infected with either WT EPEC or its ΔT3SS mutant at MOI 5:1 for 3 h. Bacterial adhesion was determined by the colony-forming assay on agar plates. Mean±SEM, n=9; ** $p<0.01$, *** $p<0.001$

Supplementary Figure 3

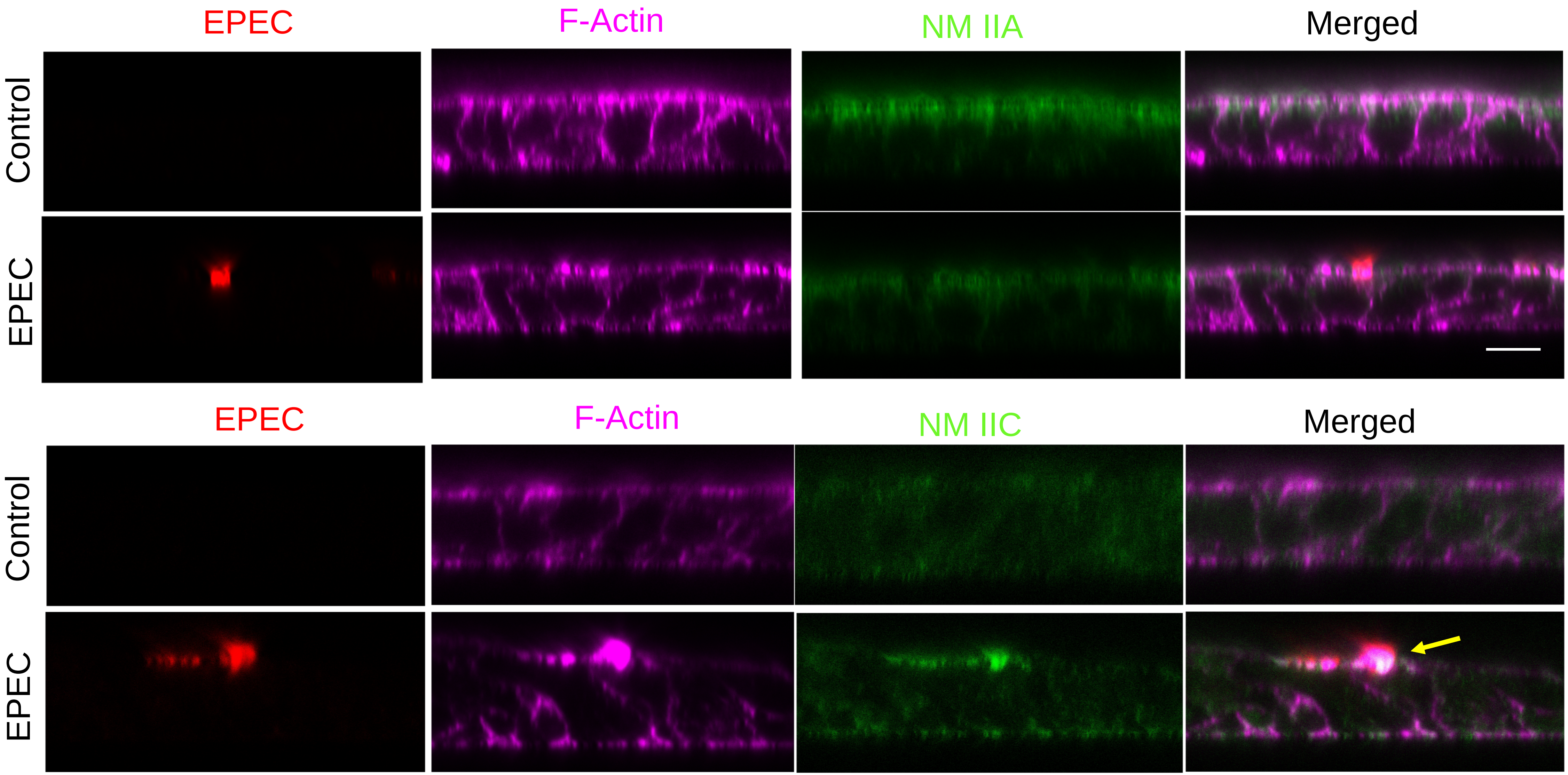


**Supplemental Figure 4: NM IIA does not accumulate at apical actin pedestals, while NM IIC is enriched at the pedestals in EPEC-infected IECs.**
Confluent, differentiated HT-29 cell monolayers were infected with EPEC at a multiplicity of infection (MOI) of 5:1 for 3 hours. Cells were then fixed and stained for F-actin (magenta), NM IIA or NM IIC (green), and EPEC (red). xz scan showing prominent apical F-actin pedestals with no detectable accumulation of NM IIA, whereas NM IIC shows clear enrichment at apical pedestals in EPEC-infected cells (arrow). Scale bar = 10 μm.

Supplementary Figure 4

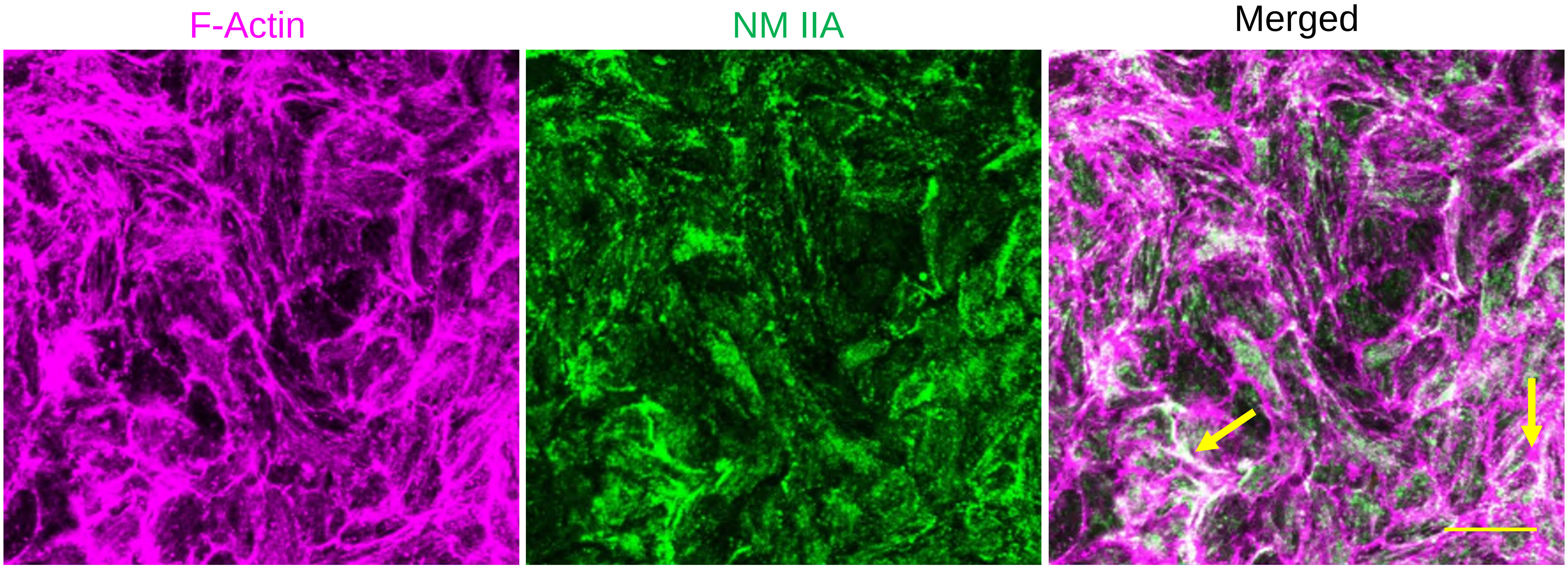


**Supplemental Figure 5: Colocalization of NM IIA with basal F-actin stress fibers in EPEC infected HT-29 epithelial cell.**
Confluent HT-29 cell monolayers were infected with EPEC for 3 h. Cells were fixed and fluorescently labeled for F-actin (magenta) and NM IIA (green). Arrows point on NM IIA enrichment at basal stress fibers. Scale bar=20 µm.

Supplementary Figure 5

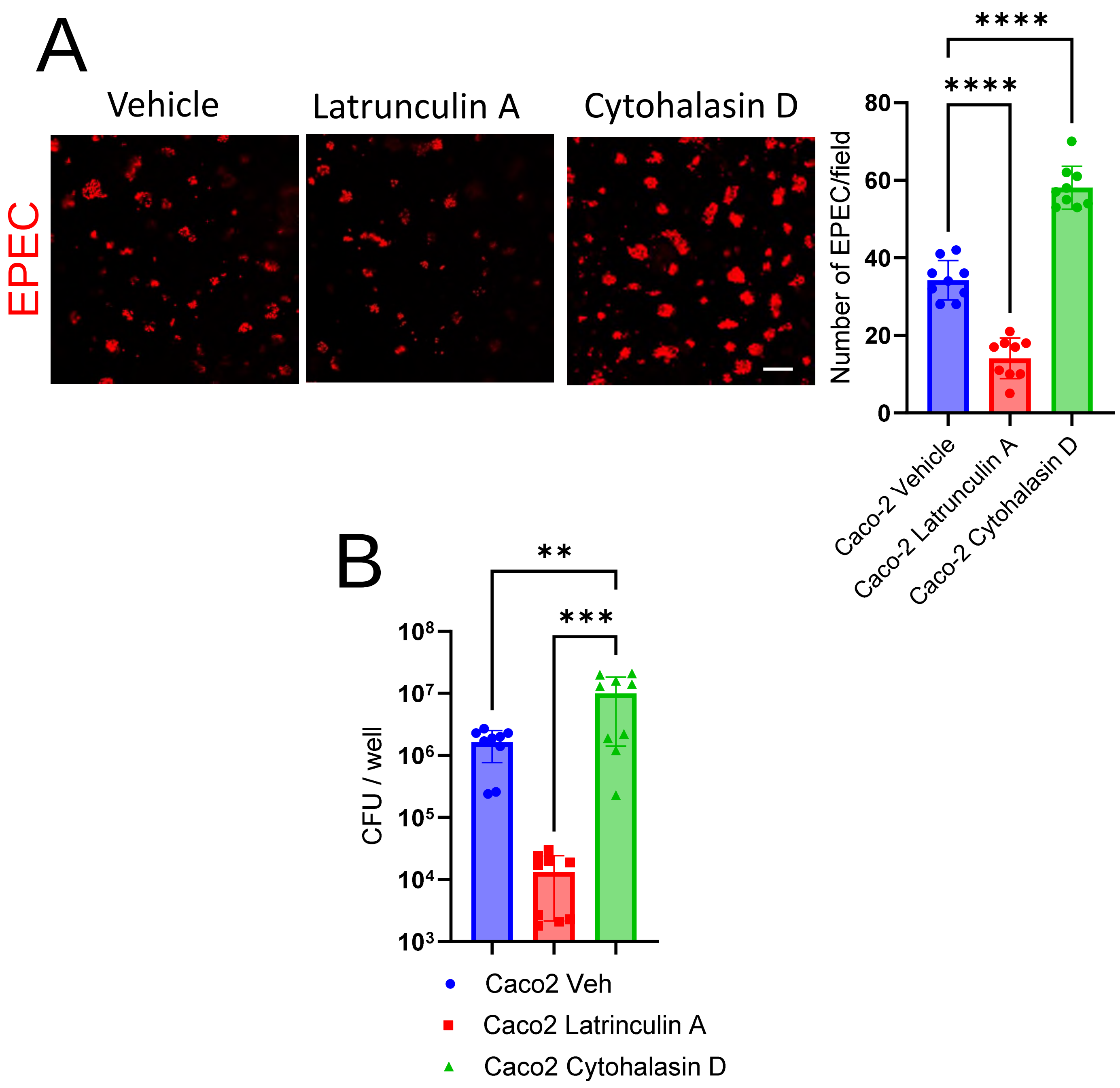


**Supplemental Figure 6: Differential effects of actin-depolymerizing drugs on EPEC attachment to IEC monolayers.**

Confluent differentiated Caco-2 BBE cell monolayers were exposed to EPEC (MOI 5:1) for 3h in the presence of either vehicle, latrunculin A (0.5 µM), or cytohalasin D (10 µM). Bacterial attachment to IEC was determined by either fluorescence labeling/confocal microscopy of LPS (**A**) representative of 3 independent experiments, or colony forming assay (**B**). Mean±SEM, n=9, **p<0.01, ***p<0.001, ****p<0.0001: scale bar 20 µm.

Supplementary. Figure 6
